# Immune-modulatory effects of low dose γ-radiation on wax moth (*Galleria mellonella*) larvae

**DOI:** 10.1101/2022.10.11.511741

**Authors:** David Copplestone, Christopher J Coates, Jenson Lim

## Abstract

Larvae of the greater wax moth *Galleria mellonella* are common pests of beehives and commercial apiaries, and in more applied settings, these insects act as alternative *in vivo* bioassays to rodents for studying microbial virulence, antibiotic development, and toxicology. In the current study, our overall aim was to assess the putative adverse effects of background gamma radiation levels on *G. mellonella*. To achieve this, we exposed larvae to low (0.014 mGy/h), medium (0.056 mGy/h), and high (1.33 mGy/h) doses of caesium-137 and measured larval pupation events, weight, faecal discharge, susceptibility to bacterial and fungal challenges, immune cell counts, activity, and viability (i.e., haemocyte encapsulation) and melanisation levels. The effects of low and medium levels of radiation were distinguishable from the highest dose rates used – the latter insects weighed the least and pupated earlier. In general, radiation exposure modulated cellular and humoral immunity over time, with larvae showing heighted encapsulation/melanisation levels at the higher dose rates but were more susceptible to bacterial (*Photorhabdus luminescens*) infection. There were few signs of radiation impacts after 7 days exposure, whereas marked changes were recorded between 14 and 28 days. Our data suggest that *G. mellonella* demonstrates plasticity at the whole organism and cellular levels when irradiated and offers insight into how such animals may cope in radiologically contaminated environments.

## INTRODUCTION

Radiation originates from natural and manmade sources and all living organisms are affected by radiation to varied extents, from cellular to whole organism levels. Humans may be exposed to low-dose radiation from medical diagnostic tools such as computed tomography (CT) scanning or high-dose radiation from radiotherapy and nuclear disasters. In humans, the response to radiation is determined by parameters including the radiation source, radiation dose (amount of radiation energy received), length of exposure, and the genetic makeup of the exposed subject (Reisz et al, 2014). The genetic and epigenetic aspects are significant and may determine the likelihood of a subject developing cancer or ability to respond to a cancer treatment, even at low doses (Ali et al, 2020). The latter is observed as some tumours either have intrinsic or acquired radioresistance through genetic mutations and selection for increased survival and proliferation (Stratton et al, 2009; Kim et al, 2015). In the environment, nuclear accidents such as the Chernobyl and Fukushima events can cause changes to wildlife, especially insects, over decades. To study the impact of ionising radiation (IR) on whole organisms, most laboratory-based experiments investigating IR effects typically adopt acute radiation exposure with conclusions relying on crude metrics such as death or sterility (Bakri et al, 2005; Dyck et al, 2021; Raines et al, 2022). From a cellular/molecular perspective, most studies on IR-induced effects on insect biological systems focus on DNA damage and its corresponding enzymatic repair processes (Costa et al. 2018). There are gaps in our understanding of IR-related health decline due to chronic exposure to background environment doses, which is between 22 – 78 nGy/h, or 0.5 - 1.0 mSv/year in the UK (Appleton and Kendall, 2022; https://www.data.gov.uk/dataset/568e58c0-6404-4a8a-9654-4440245fb6e4/ambient-gamma-radiation-dose-rates-across-the-uk). Low doses of gamma radiation result in a small amount of damage and repair which might lead to cell necrosis/death and has the potential of driving childhood cancers though in some cases, organisms adopt resistance to low levels of radiation due to selection pressures, a process not completely understood due to a lack of replicative studies (Cuttler 2007; Mazzei-Abba et al., 2020; Little et al., 2010). However, and more importantly, IR impacts to organisms go beyond DNA damage, altering expression of genes associated with immunity, though its contribution remains unclear (Lumniczky et al, 2021). Understanding this process can offer alternative biomarkers to determine whether IR is having any substantial biological (or specifically, immunological) impact (Bednarski and Sleckman 2019). All this is crucial to our understanding of contaminated natural environments such as the Chernobyl and Fukushima, and their long-term management and environmental restoration.

Therefore, to understand the role chronic low-dose radiation exposure has on the immune system, we used insect larvae from the Greater wax moth, *Galleria mellonella*. Wax moth larvae are much larger than traditional insect models like Drosophilids, thereby making it easier to obtain large number of cells for downstream experiments, and no specialist equipment and training are needed (Coates et al., 2019; Emery et al., 2019, 2021). They are also ethically more acceptable than vertebrate work therefore allowing larger experimental numbers to be utilised and thus improving statistical discrimination (Mather 2001; Mylonakis 2008; Lionakis 2011). They are amenable to physiological manipulations in their life history and ecology (Rolff & Siva-Jothy, 2003). A key biological advantage is that *G. mellonella* possesses innate immunity, without adaptive immunity, thereby simplifying this study. Like mammalian innate immunity, there are both cellular and humoral components. The cellular component comprises a heterogenous population of circulating haemocytes that are highly similar to human and mouse neutrophil/macrophage counterparts (Browne et al 2013). They play a critical role in pathogen phagocytosis, encapsulation, nodulation, degranulation, etosis, and hemostatic events (Kay et al 2019; Krachler et al, 2021). The humoral defences are comprised of the soluble immune factors of the haemolymph, which includes activation of the pro-phenoloxidase cascade, production of the antimicrobial pigment melanin and the darkening of the larval integument (Tsai et al 2016; Whitten and Coates, 2017).

Herein we show that while gamma radiation from a caesium-137 source had little effect on the weight of *G. mellonella* larvae over time, it impacted the volume of faecal discharge and rate of pupation. Immune responses to IR varied, with an increase in encapsulation, susceptibility to infection and fungal load. On a cellular level, endocytosis, melanisation, and lysosomal membrane instability increased with IR. Together, these data provide a basis for further studies investigating how IR and other non-gamma radiation sources can influence the immune system in a tractable experimental system that can be extrapolated to radiologically contaminated sites.

## MATERIALS AND METHODS

### Reagents

All key reagents, such as Grace’s Insect medium (G9771), alkaline phosphatase (10713023001) and Lysogeny Broth (L3522), were purchased from Sigma-Aldrich, UK (now MERCK) in their purest form. Heat-inactivated foetal bovine serum (FBS), yeast extract-peptone-dextrose (YPD), phosphate buffered saline (PBS, pH 7.4) tablets, 8-Hydroxy-1, 3, 6-pyrenetrisulfonic acid trisodium salt (HPTS), neutral red and thiazolyl blue tetrazolium bromide (MTT) were from Fisher Scientific, UK.

### *Galleria mellonella* maintenance

Final instar larvae of the greater wax moth, *G. mellonella*, were sourced from Livefoods Direct (UK) and stored in wood shavings in the dark at 15°C. Healthy larvae weighing between 0.2 and 0.4 g were used in all experiments, i.e., displaying no visual signs of damage, infection, or melanisation. Experimental manipulation of insects was approved by the University of Stirling’s Animal, Welfare and Ethical Review Body (AWERB (16/17) 17 New Non-ASPA).

### Exposure of *Galleria mellonella* larvae to ionising radiation

The radiation facility at the University of Stirling (Scotland, UK) was used for all exposure experiments – temperature controlled (17°C or 25°C) with a caesium-137 gamma emitting source. A gamma-emitting source was selected because most invertebrates in the Chernobyl Exclusion Zone (CEZ) are exposed to such radiation (Beresford et al 2020). Shelves were placed at 1 m (“High”), 4.5 m (“Medium”) and 9 m (“Low”) from the source for the samples. Shelves were also placed in a shielded area within the facility and designated as “control” (**Figure 1**). We confirmed the doses using dosimeters (Model 23-1 Electronic Personal Dosimeter, Ludlum Measurements Inc., USA) to take readings around the facility with background radiation reading at the University of Stirling at 0.11±0.01 μGy/h (Burrows et al, 2022; Raines et al, 2020; Goodman et al, 2019). Larvae were distributed around the facility via four groups (“Control”, “Low”, “Medium” and “High”), which was reflective of the different dose rates received (**Table 1**) and are comparable with those found in the CEZ (up to 250 μGy/h; Beresford et al 2020). At specified times (7 – 42 days), larvae were taken out for analysis.

**Figure 1.**
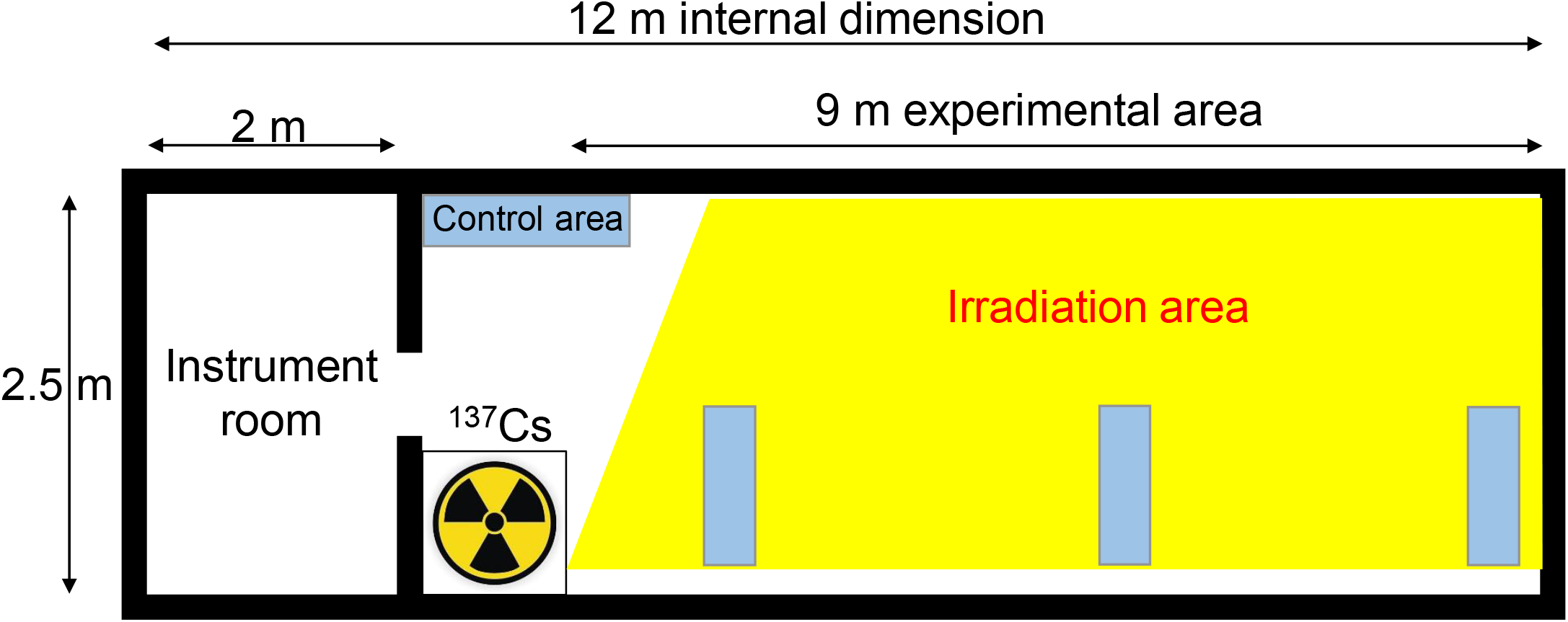
Schematic diagram of the radiation facility with a ^137^Cs source. Shelving units (blue boxes) were placed in the path of the radiation source, as well as in a zone outside the radiation area (control area).

**Table 1:** Averaged dose rates in the radiation facility (mGy / hr). N defines the number of readings taken over a 2-year period.

|  | N | Mean | SD |
| --- | --- | --- | --- |
| Control | 10 | $4.11 \times 10^{-4}$ | $1.77 \times 10^{-5}$ |
| Low | 16 | 0.014 | 0.001 |
| Medium | 11 | 0.056 | 0.007 |
| High | 18 | 1.333 | 0.253 |

### Bacterial cell growth conditions

Single colonies of *Photorhabdus luminescens* subsp. laumondii TT01 (NCIMB 14339) were cultured in Lysogeny Broth (LB) broth for 16 h at 30°C in an orbital incubator at 220 rpm. Cells were centrifuged at 6000 x *g* for 2 min at room temperature, washed three times in PBS (10 mM phosphate buffer, 2.7 mM KCl, 137 mM NaCl, pH 7.4) and optical density (OD) at 600 nm was measured with a cell density meter (WPA, Biochrom). Bacterial cells were serially diluted in PBS to an OD of 0.00001, which represents ~8 x10^3^ colony forming units (CFUs) per ml prior to experimental use.

### Fungal cell growth conditions

*Candida albicans* SC5314 (kindly donated by Dr Rebecca Hall, Kent, UK), *Cryptococcus neoformans* and *C. gattii* (kindly donated by Prof Robin May, Birmingham, UK) were grown to the stationary phase in liquid YPD medium for 24 h at 25 °C on a rotator at 20 rpm (or 37°C, 200 rpm for *C. albicans*) as shown previously (Lim et al, 2018). Yeast cells were centrifuged at 3,000 x g for 2.5 min, washed three times in PBS, and counted with a haemocytometer prior to use.

### *Photorhabdus luminescens* infection assay

Larvae weighing at least 0.2 g were injected with 20 μl of PBS or 20 μl of a bacterial suspension containing 8 × 10^7^ CFUs/ml (PBS) of *P. luminescens*, i.e., 1.6 x10^6^ per larva. Larvae were inoculated by intrahaemocoelic injection into the last right pro-leg using a 25-gauge hypodermic needle (BD Microlance™) attached to a 1 ml syringe (BD Plastipak™). Treated larvae (n = 30 per condition over 3 independent experiments) were incubated at 30°C and scored for mortality every 24 h.

### Fungal load assay

This protocol was adapted from Mowlds et al (2010). As before, larvae were injected with a low dose (1 × 10^6^ / larvae) of either *Cryptococcus neoformans* H99, *C. gattii* R265 or *Candida albicans* SC5314 and incubated at 25 °C for 16 h. Larvae were homogenized in 3 ml of sterile PBS, before being serially diluted with PBS. Approximately 100 μl of the resulting dilutants were plated on LB agar containing erythromycin (100 μg/ml) to prevent bacterial growth. Plates were incubated at 25 °C for 16 h prior to enumeration of colony forming units. Data are expressed as CFUs per larva.

### Total haemocyte counts using trypan blue exclusion

Haemolymph was extracted, pooled and mixed 1:1 with 0.4% trypan blue dissolved in PBS. Total viable haemocyte numbers were determined by dye exclusion using a haemocytometer (FastRead) and adjusted to 1 × 10^6^ cells per 1 ml PBS for subsequent *in vitro* experiments.

### *In vivo* melanisation determination

This protocol was adapted from Kloezen et al (2015). Four dishes with larvae were distributed into the 4 different parts (“Control”, “Low”, “Medium” and “High”) of the radiation facility at 17 °C to slow down the process of pupation and limit the scope of experiments to the larvae only. Between 3 and 5 representative larvae per condition were randomly selected at defined time intervals under radiation conditions. Haemolymph was extracted through an incision behind the head of the larvae. For melanisation, approximately 20 μl haemolymph per larva was added to 20 μl ice-cold PBS containing 0.37% β-mercaptoethanol and kept on ice to stop further increases in melanin concentration. Samples were transferred to a 96-well U-bottom plate and absorbance at 405 nm was measured using a Molecular Devices VersaMax plate reader. Sample sizes included nine larvae per condition across three independent experiments.

### *In vivo* encapsulation assay

Four dishes with larvae were distributed into the 4 different parts (“Control”, “Low”, “Medium” and “High”) of the radiation facility at 25 °C and left for 7 days. Larvae were surface sterilised with 70% ethanol and a 3 mm piece of nylon thread (0.45 mm diameter, 100% polyamide, efco creative GmbH) was inserted gently into the haemocoelic cavity parallel to the GI tract via its last right pro-leg using a pair of fine forceps. Larvae were incubated for 24 h at 30 °C, before frozen at −20 °C for further analysis. Larvae were defrosted and dissected to retrieve the nylon implant. Implants were imaged – at both ends – using an upright (light) microscope (CX31, Olympus) with an eyepiece camera (BF960, Swift Optical Instruments Ltd). Percentage of nylon thread covered by capsule formation was determined by measuring capsular area by using the ImageJ software (National Institutes of Health) and related to the entire thread as visualised. Sample sizes included 12 larvae per condition across two independent experiments.

### *In vivo* faecal bacterial cell viability assay

Four dishes with larvae were distributed into the 4 different parts (“Control”, “Low”, “Medium” and “High”) of the radiation facility at 25 °C and left for 7 days. Faecal material was collected, weighed, and resuspended in sterile distilled water, shaking at 1000 rpm for 30 min, before adjusted to 10 mg/ml. After centrifugation at 16 162 xg (Sigma 1-14, Sigma Laborzentrifugen GmbH) for 5 min, faecal material was resuspended with water. Approximately, 90 μl of faecal suspension was mixed with 10 μl (10x) buffer and 0.5 μl (0.5 U) calf intestinal phosphatase (APMB-RO, Roche, Sigma-Aldrich) and incubated at 37 °C for 60 min. 100 μl of ATP-depleted faecal suspension was mixed with 100 μl BacTiter-Glo™ Microbial Cell Viability Assay (Promega) in a clear, flat-bottomed 96-well plate (Greiner) and left on a platform shaker at 25 rpm for 5 min at room temperature. Plates were read for luminescence using the GloMax system (Promega) with pre-installed settings.

### *In vivo* adhesion assay

This protocol was adapted from Humphries (2009). As before, 3-5 representative irradiated larvae at 17 °C were removed from the facility at defined time intervals. Haemolymph was extracted, pooled and total haemocyte counts determined by trypan blue exclusion as described earlier. Approximately, 1 × 10^5^ haemocytes per well were seeded into a 96-well plate and left to adhere for 20 min at room temperature. Nonadherent cells were washed three times using PBS and attached cells were fixed using 100 μl 5% (w/v) glutaraldehyde for 20 min at room temperature. Wells were washed 3 times with distilled water prior to staining with 100 μl 0.1% (w/v) crystal violet in 200mM MES, pH 6.0, for 1 h at room temperature. Wells were washed 3 times with distilled water before dye was solubilised with 100 μl 10% (v/v) acetic acid and agitated on an orbital shaker at 150 rpm for 5 min at room temperature. Absorbance was measured at 570 nm using a plate reader (VersaMax, Molecular Devices).

### *In vivo* neutral red retention assay

This protocol was adapted from Hu W et al (2015). As before, 3-5 representative irradiated larvae at 17 °C to slow down the process of pupation were removed from the facility after a specific amount of time. Haemolymph was extracted and pooled and its total haemocyte count determined by trypan blue exclusion as described earlier. Approximately 1 × 10^5^ haemocytes were seeded into a 96-well plate and left to adhere for 60 min in Grace’s medium supplemented with 0.35g/L NaHCO_3_, pH 6.2 at room temperature. Neutral red dye solution (200 μM in Grace‘s medium) was added and samples were left in the dark for 90 min for dye uptake. Wells were washed with fixative solution (1% formaldehyde, 1% calcium chloride) for 2 min before the addition of an extraction buffer (1% acetic acid, 50% ethanol) and left in the dark for 20 min at room temperature. Absorbance of extracted dye was measured using a microplate reader (VersaMax, Molecular Devices) at a wavelength of 570 nm.

### *In vitro* assays

Pooled haemolymph from 4-10 larvae was counted and 100,000 haemocytes per well distributed into 96-well plates, in Grace’s medium. Plates were placed in the 4 different locations (“Control”, “Low”, “Medium” and “High”) of the radiation facility and irradiated for 48 h. For quantitation of **melanisation**, wells containing haemocytes were washed with ice-cold PBS containing 0.37% β-mercaptoethanol and absorbance measured at 405 nm using a VersaMax (Molecular Device) microplate reader. For **cell viability**, cells in wells were washed in PBS and 0.5mg/ml of MTT (in Grace+ medium) added and incubated for 2-4 h at 30 °C. Medium was removed, cells washed in PBS and solubilised and shaken with DMSO. Plate was read on microplate reader (VersaMax, Molecular Devices) at 595 nm. For **pinocytosis**, cells in wells were washed in PBS and 5 mg/ml of HPTS dye (in PBS) added and incubated with gentle rocking at 30 °C for 120 min. After washing in PBS at least 4 times to remove all extracellular dye, samples were fixed using 4% PFA/PBS for 10 minutes. Samples were transferred into a plate reader and fluorescence was recorded with excitation and emission filters set at wavelengths 405 nm and 520 nm, respectively. To quantify cellular uptake, fluorescence readings were divided by the average value obtained from the corresponding viability (MTT) assays (fluorescence/absorbance). All determinations were performed on at least three independent experiments.

### Statistical analyses

We used log-rank (curve comparison) tests to assess survival data from bacterial and pupation levels, and linear regression for larval weight. We applied ANOVA followed by Tukey’s multiple comparisons test to assess the effects of radiation dose and length of exposure on all remaining experimental endpoints. All analyses were performed in GraphPad Prism 9.3.1., San Diego, California USA, www.graphpad.com. Samples sizes can be found within the respective methods sections.

## RESULTS

### Whole organism responses to radiation

*Galleria mellonella* larvae decreased in weight over 42 days (grams, 17 °C), regardless of the radiation exposure rates (low, medium, high dose = 14.11, 56.45, 1343.66 mGy after 42 days). However, while the rates of weight loss differed significantly over time from each other (as determined by simple linear regression of weights; F = 3.656. *p* = 0.0134), only insects subjected to the medium radiation dose were significantly different from the control (F = 4.224, *p* = 0.0424; **Figure 2A**) even though there were no significant differences in weight at the end of experiment on day 42 (**Figure 2B**), with little difference in rates of pupation and death at this temperature (**Supplementary Figure 1**). Interestingly, the rate of weight loss for a high dose of radiation (*p* = 0.02; D’Agostino-Pearson) is very similar to the control (*p* = 0.005; D’Agostino-Pearson) which suggests an inherent radioprotection or better recovery from the impact of radiation. Interestingly, at higher temperature (25 °C), pupation was faster with larvae exposed to a higher dose of radiation (**Figure 3**, χ^2^ = 86.76, df = 3, *p* < 0.0001).

**Figure 2.**
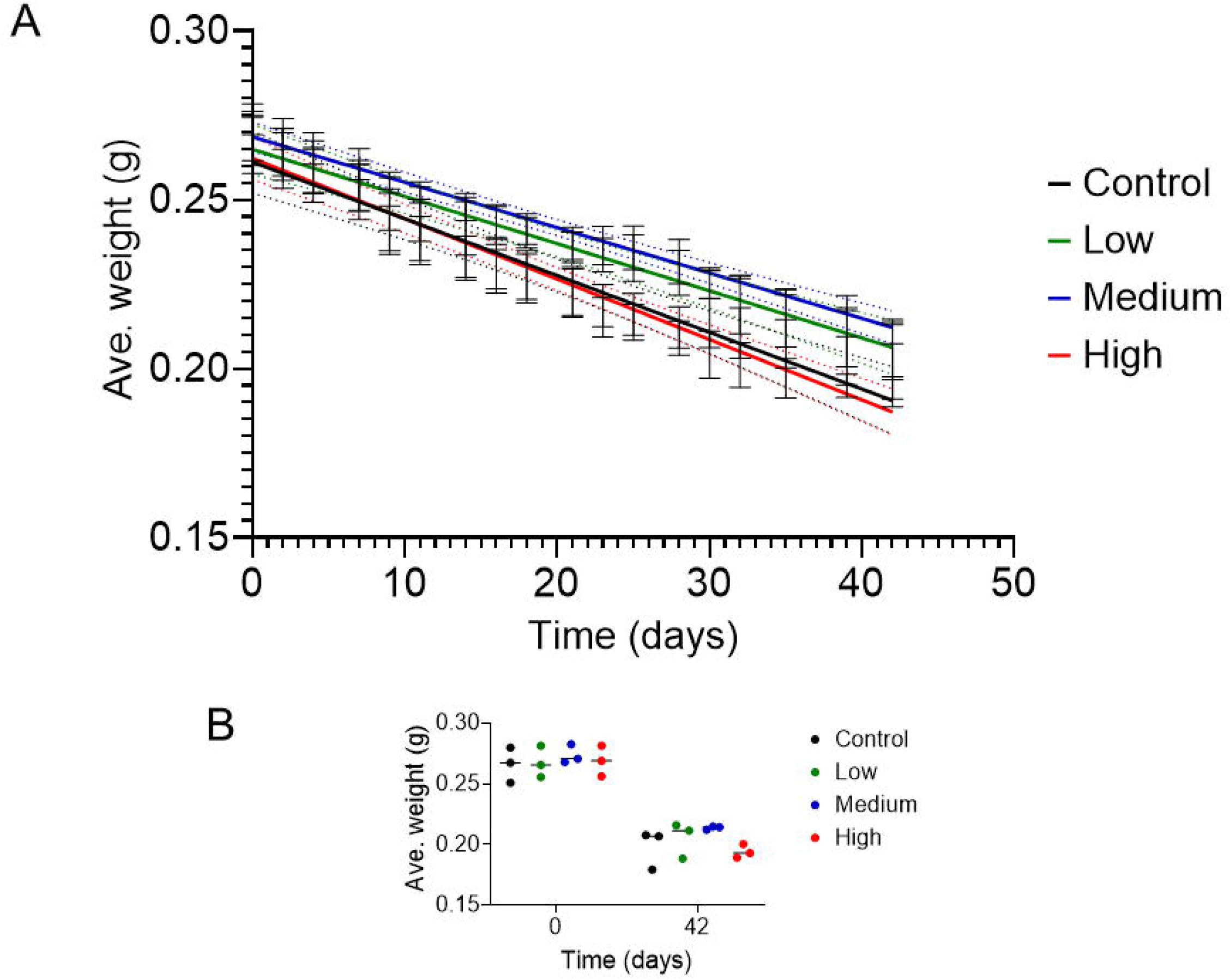
Ionising radiation had little impact on average weights (grams) of *Galleria mellonella* larvae. *G. mellonella* larvae were exposed to variable dose rates of ionising radiation at 17 °C for the indicated period of time. Larvae were weighed daily and plotted as a connecting line graph with SD bars (a) and as an interleaved scatter plot of the start and end of the experiments with SD bars (b). Results were based on the average of three independent experiments (n = 30).

**Figure 3.**
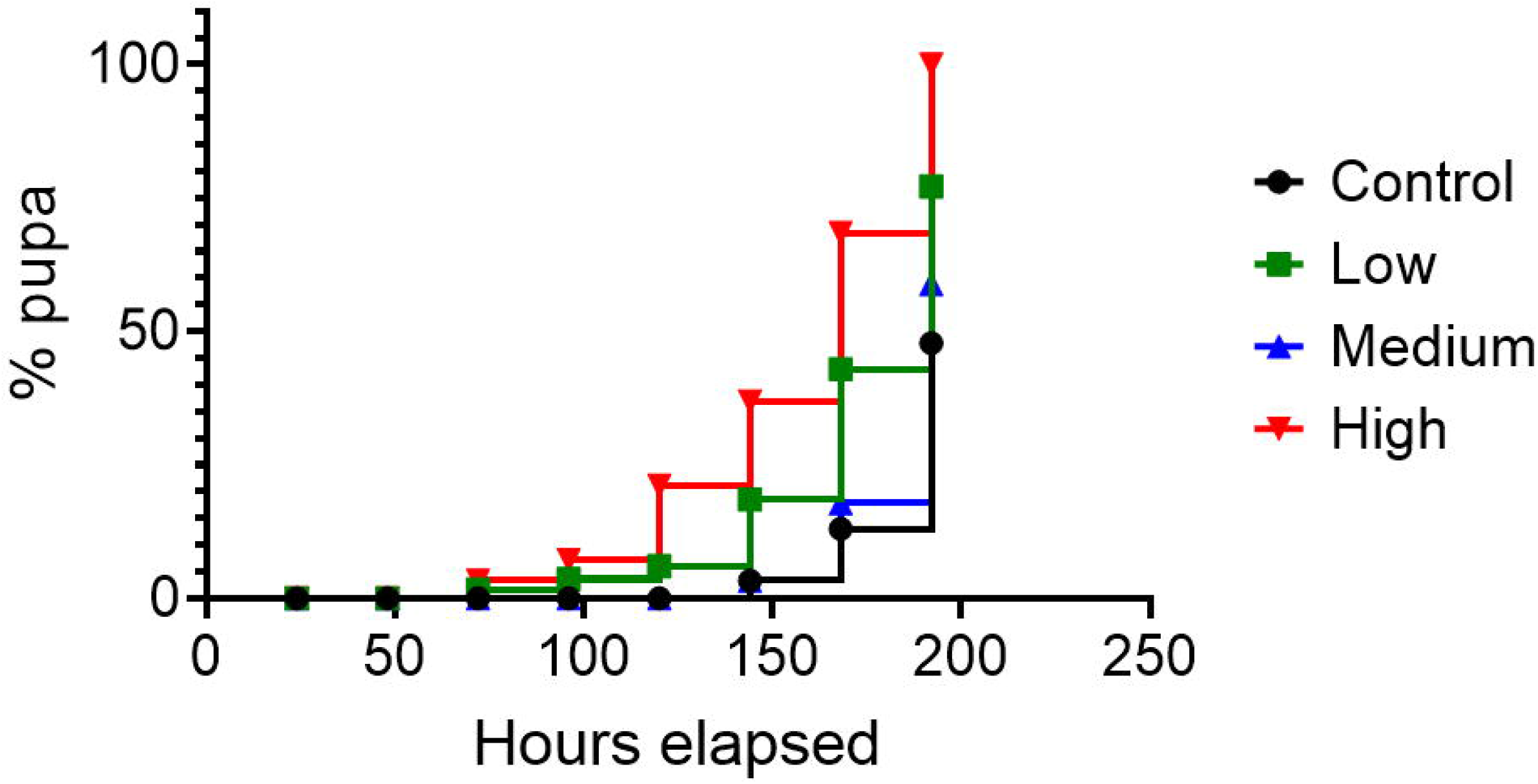
Ionising radiation affected pupation of *Galleria mellonella. G. mellonella* larvae were exposed to variable dose rates of ionising radiation at 25 °C for the indicated period of time. Larvae were checked daily for pupa formation. Results were based on the total of two independent experiments (n = 24).

Moreover, weight of excrement (i.e., faecal load) produced by each larva after radiation exposure was investigated with visual discrepancies were observed. The amount of faeces did not change much over time in control and low dose radiation conditions. However, there were significant decreases in faeces produced by larvae exposed to medium and high dose of radiation over time (**Figure 4A**; **Medium** at Day 0, 0.0048±0.00008 g c.f. Day 21, 0.0035±0.00026 g and **High** at Day 0, 0.0050±0.00004 g c.f. Day 21, 0.0036±0.00018 g). Interestingly, there was a subtle higher amount of faeces from larvae exposed to a medium and high dose of radiation compared to the control at day 7 (0.004±0.0002 (**control**) c.f. 0.005±0.0001; *p* = 0.01 (**medium**) c.f. 0.005±0.0002; *p* = 0.59 (**high**)). This could be due to the initial levels of environmental stress placed on the larvae. Interestingly, we found higher levels of viable bacteria from the faecal materials obtained from larvae exposed to medium dose of radiation compared to control after 7 days (**Figure 4B;** 36620±13369 RLU (**control**) c.f. 444251±111079 RLU (**medium**); *p* = 0.001). This suggests major effects of radiation on the gut and its flora occurs in the first 7 days, before they adapt to the environmental stressor.

**Figure 4.**
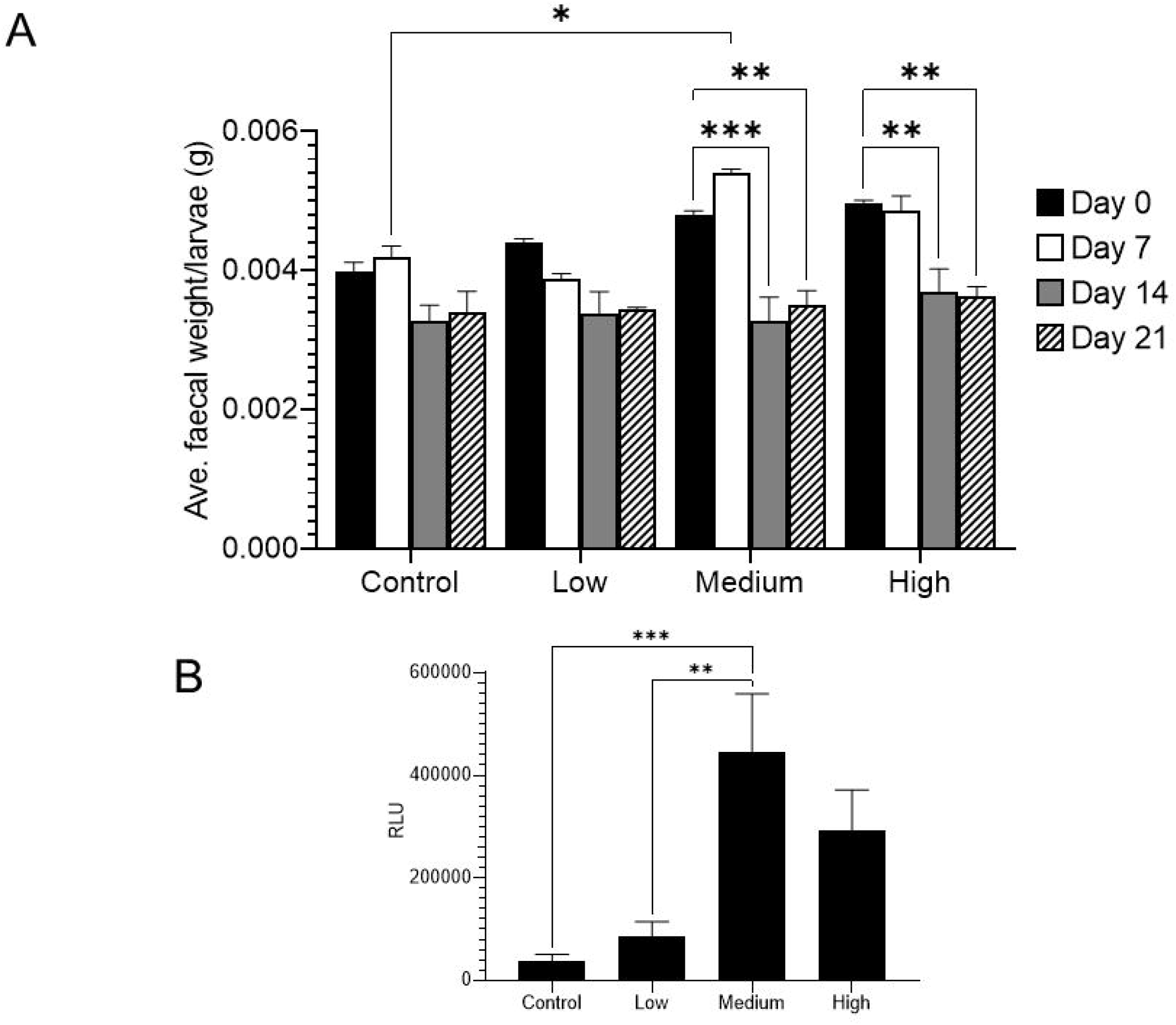
Radiation affected faecal discharge from *Galleria mellonella. G. mellonella* larvae were exposed to gamma radiation from a Cs-137 source for up to 21 (A) or 7 (B) days and faecal material was weighed (grams) (A), resuspended with water and tested for viable bacteria (B), as described in the Materials and Methods. Significance was determined by two- (A) and one- (B) way ANOVA, and a Tukey’s multiple comparisons test. (***) p ≤ 0.001, (**) p ≤ 0.01, (*) p ≤ 0.05. Results are expressed as the mean ± SD of at least two independent experiments (n = 36/sample).

### Radiation impact on larval susceptibility to infection

Larvae were exposed to radiation at different intensities before receiving an intrahaemocoelic dose of *Photorhabdus luminescens* via injection. This entomopathogenic bacterium is known to infect *G. mellonella* and other lepidopteran species (Shahina et al 2011; Wu et al 2014; Patterson et al 2015). Short (≤14 days) exposure times to radiation of different doses did not affect larval susceptibility to *P. luminescens* (**Figure 5**). However, longer (21 days) exposure times appear to negatively affect larvae susceptibility to *P. luminescens* infection, significantly at high doses when compared to the control (*p* = 0.001), marginally for medium (*p* = 0.023) doses but not for low (*p* = 0.06) doses (**Figure 5**).

**Figure 5.**
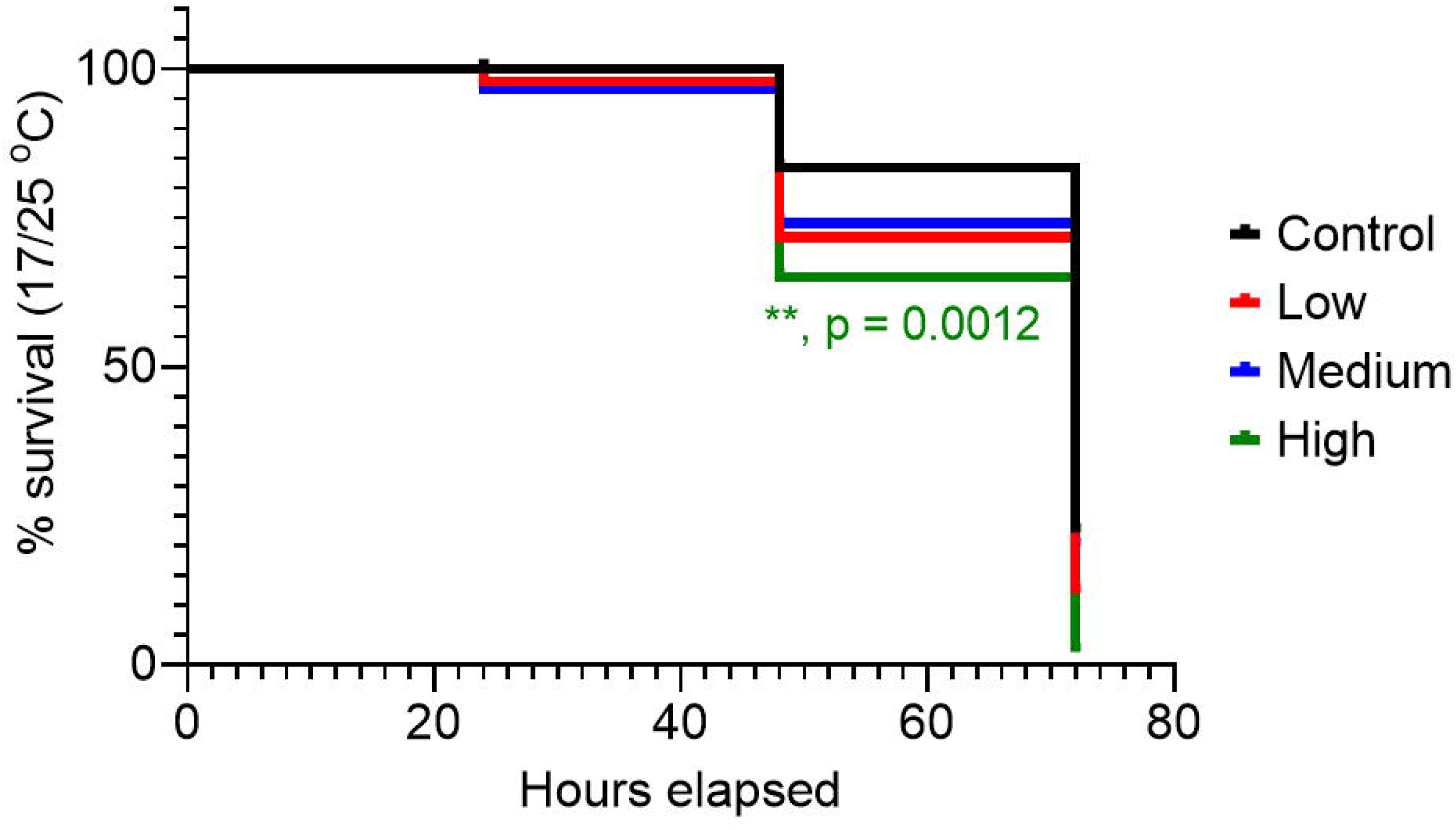
Long-term exposure of *Galleria mellonella* to ionising radiation leads to susceptibility to *Photorhabdus luminescens* infection. *G. mellonella* larvae were irradiated for 21 days at 17 °C before injected with 20 μl of a suspension containing 8 × 10^7^ CFU/ml of *P. luminescens* bacteria. Larvae were incubated at 25 °C and monitored daily for survivability. Data were means ± s.e.m. Significance in survival of larvae pre-exposed to radiation compared to non-radiated larvae was determined using a log-rank (Mantel-Cox) test (n = 30/sample).

To gain further understanding, we explored the capacity of the insect host to prevent replication of microbes. To achieve this, we determined the fungal load (i.e., the number of yeast cells per larva) of pre-radiated larvae injected with yeasts. These results demonstrate that, for control (non-radiation) conditions, following an initial inoculum of 100 × 10^4^ yeast cells per larva, the fungal load increased to 189±14.5 × 10^4^, 186.5±9.5 × 10^4^ and 271.5±36.5 × 10^4^ per larva in 16 hours post inoculation for *C. albicans, C. neoformans* and *C. gattii*, respectively. However, under low dose radiation conditions, the fungal load for *C. albicans* and *C. gattii* decreased to 84.5±3.5 × 10^4^ (*p* = 0.02) and 144±0 × 10^4^ (*p* = 0.005) per larva in 16 h, respectively. Interestingly, at higher radiation dose, fungal load for *C. gattii* increased to 386±20 × 10^4^ (*p* = 0.01) per larva in 16 h (**Figure 6**). This suggests the immune system of *G. mellonella* was “primed” at low radiation dose, thereby allowing partial clearance of some of the yeast strains but chronic exposure of radiation has an adverse effect on *G. mellonella*..

**Figure 6.**
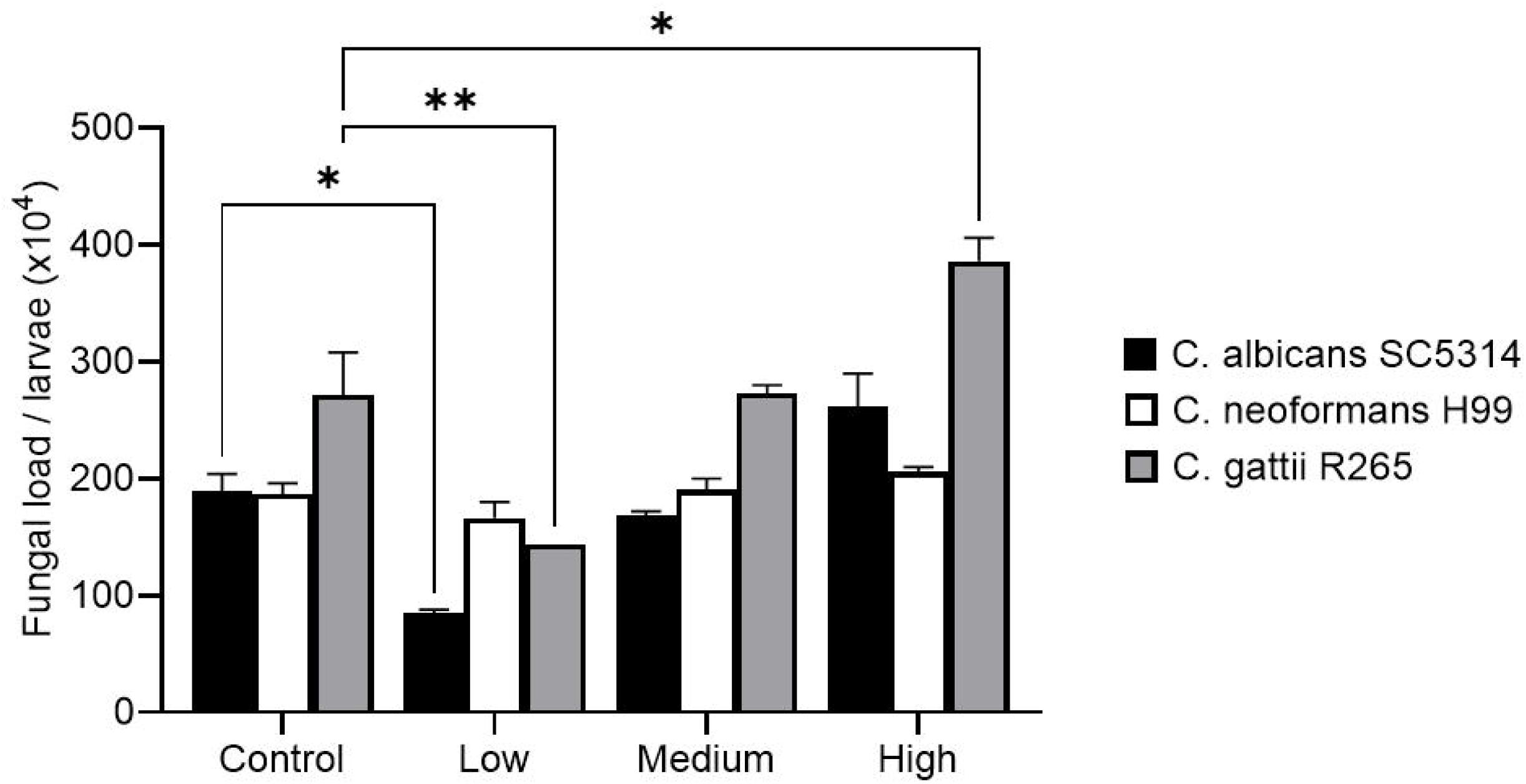
Impact of radiation on fungal load in *Galleria mellonella. G. mellonella* larvae were irradiated for 7 days prior to an intrahaemocoel challenge with 1×10^6^ *Candida albicans* strain SC5314, *Cryptococcus neoformans* strain H99 and *Cryptococcus gattii* R265. and incubated at 25 °C for 16 h. The fungal load was ascertained by serially diluting homogenized larvae and plating aliquots onto erythromycin containing LB agar plates. Results are expressed as the mean ± SEM of two independent experiments (n = 6/sample). A significant change in fungal load per larva relative to that in the PBS treated larvae at p < 0.05 and p < 0.005 is indicated by * and **, respectively.

### Radiation affects encapsulation and melanin production

To further characterise the larval immune response to chronic radiation, we quantified the amount of immune cell (haemocyte) driven encapsulation of a nylon implant inserted into the haemocoel of *G. mellonella* larvae. One of the early and effective defence mechanisms against pathogens (or any foreign objects) is encapsulation, which is followed by the release of pro-phenoloxidase (PPO) and the subsequent production of melanin, a process called melanotic encapsulation (Nappi and Vass 1993; Strand and Pech 1995; Whitten and Coates 2017). We found only larvae exposed 7 days to high radiation (29.18±6.35; *p* = 0.04) had significantly more capsular material around the thread compared to its low dose (16.11±3.4) (**Figure 7**).

**Figure 7:**
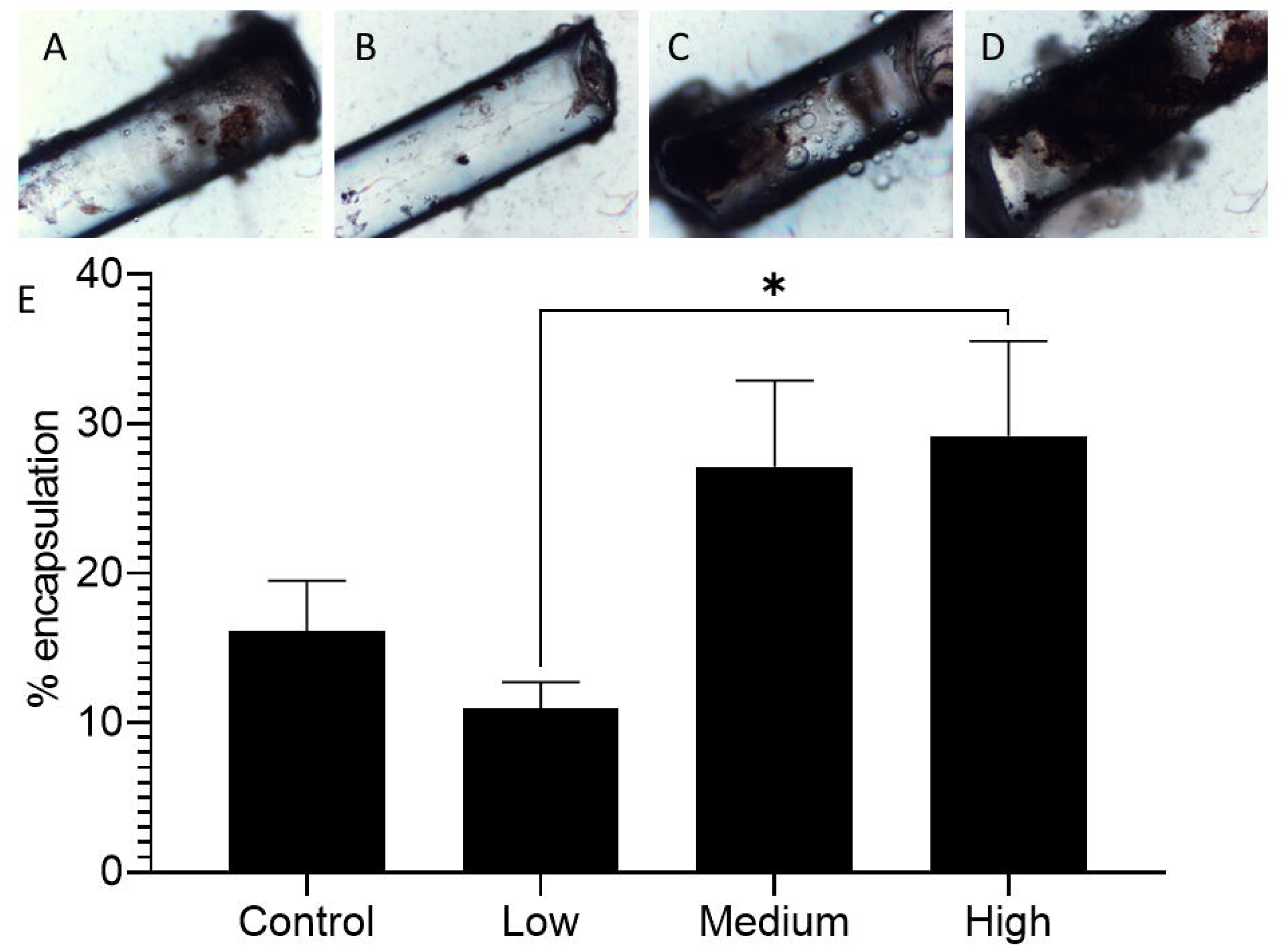
Ionising radiation affected encapsulation in *Galleria mellonella. G. mellonella* larvae were irradiated for 7 days at 25 °C before a piece of nylon thread inserted in the haemocoelic cavity. Larvae were incubated at 25 °C for 24 hand frozen. Then larvae were dissected and nylon thread removed and imaged under light microscopy. % encapsulation was the surface area of thread covered by capsule material. Data are means ± s.e.m. Significance compared to irradiated larvae with non-radiated larvae was determined by one-way ANOVA, and Tukey’s multiple comparisons test. (*) p ≤ 0.05, otherwise not significant.

Next, we wanted to determine if the changes in encapsulation and melanin production were due to changes with viable haemocytes. Total viable haemocyte count (THC) was determined using trypan blue exclusion assay. While there were no significant differences in THC in the first 7 days of exposure to radiation, differences were seen after that. At days 14 and 21, THC decreased at low (e.g. day 21, 285±49.5 × 10^4^ cells/ml) and medium (e.g. day 21, 2355±49.5 × 10^4^ cells/ml) radiation doses compared to the control (e.g. day 21, 405±49.5 × 10^4^ cells/ml), but maximal at high dose radiation (985±77.8 × 10^4^ cells/ml). Interestingly, at Day 28, THC was maximal at medium (1185±35.4 × 10^4^ cells/ml) radiation dose but not at low (690±42.4 × 10^4^ cells/ml) and high (372±24.7 × 10^4^ cells/ml) radiation doses, when compared to the control (540±42.4 × 10^4^ cells/ml) (**Figure 8A**).

**Figure 8.**
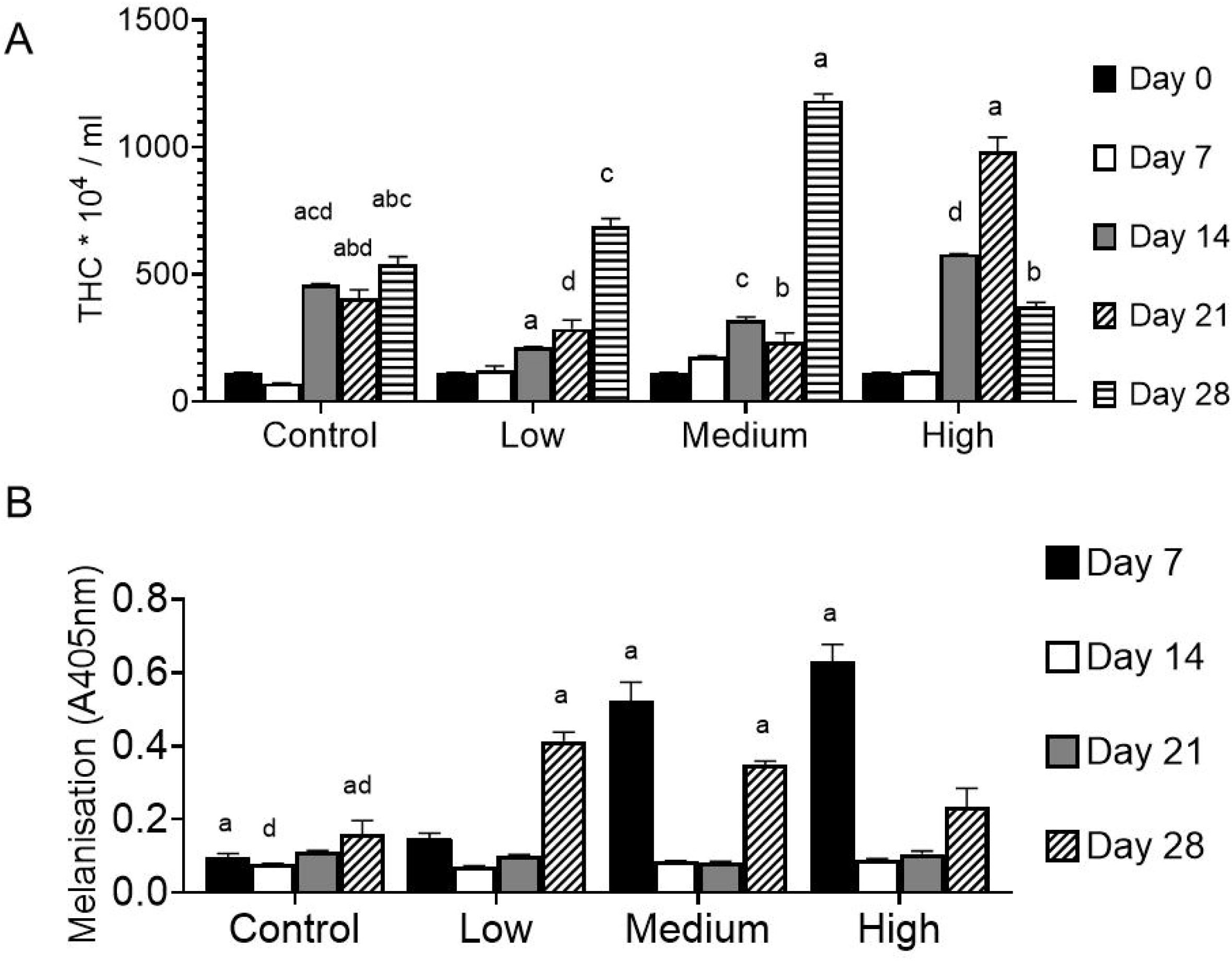
Radiation affected total haemocyte counts and melanin production in *Galleria mellonella. G. mellonella* larvae were exposed to gamma radiation from a Cs-137 source for up to 21 days and at the indicated times, pooled haemolymph was counted for haemocytes microscopically **(A)**, or measured for melanin, spectrophotometrically **(B)**. Significance was determined by two-way ANOVA, and a Tukey’s multiple comparisons test. (***) p ≤ 0.001, (**) p ≤ 0.01, (*) p ≤ 0.05. Results are expressed as the mean ± SD of at least two independent experiments (n = 36).

The encapsulation data complemented with melanin production and at day 7, melanin was produced in a (radiation) dose-dependent manner suggesting a potential protective role that melanin possess (**Figure 7** c.f. **Figure 8B**). However, after 28 days of irradiation, melanin was produced maximally under low and medium radiation (*p* = 0.13) dose conditions (**Figure 8B**). This suggests a priming effect of low dose radiation to maintain a baseline level of protection for the insect larvae.

### Radiation affects lysosomal integrity and adhesion properties of cells

To determine what other cellular functions are perturbed by ionising radiation, two further *in vivo* assays were performed, namely the adhesion assay and the neutral red retention (NRR) assay to determine cell adhesive properties and lysosomal membrane integrity, respectively. *G. mellonella* larvae was exposed to ionising radiation and haemolymph pooled and tested for lysosomal membrane stability by measuring NRR at 90 min time points. Results were analysed and statistically compared to the control group (**Figure 9A**). Lysosomal membrane stability showed a significant decrease upon 21-day exposure to high dose of radiation compared to control (0.147±0.004 vs. 0.077±0.016, *p* = 0.007) suggesting toxic effects on lysosomes due to radiation. However, no significant effects were observed on at other dose conditions, suggesting lower toxicity.

**Figure 9.**
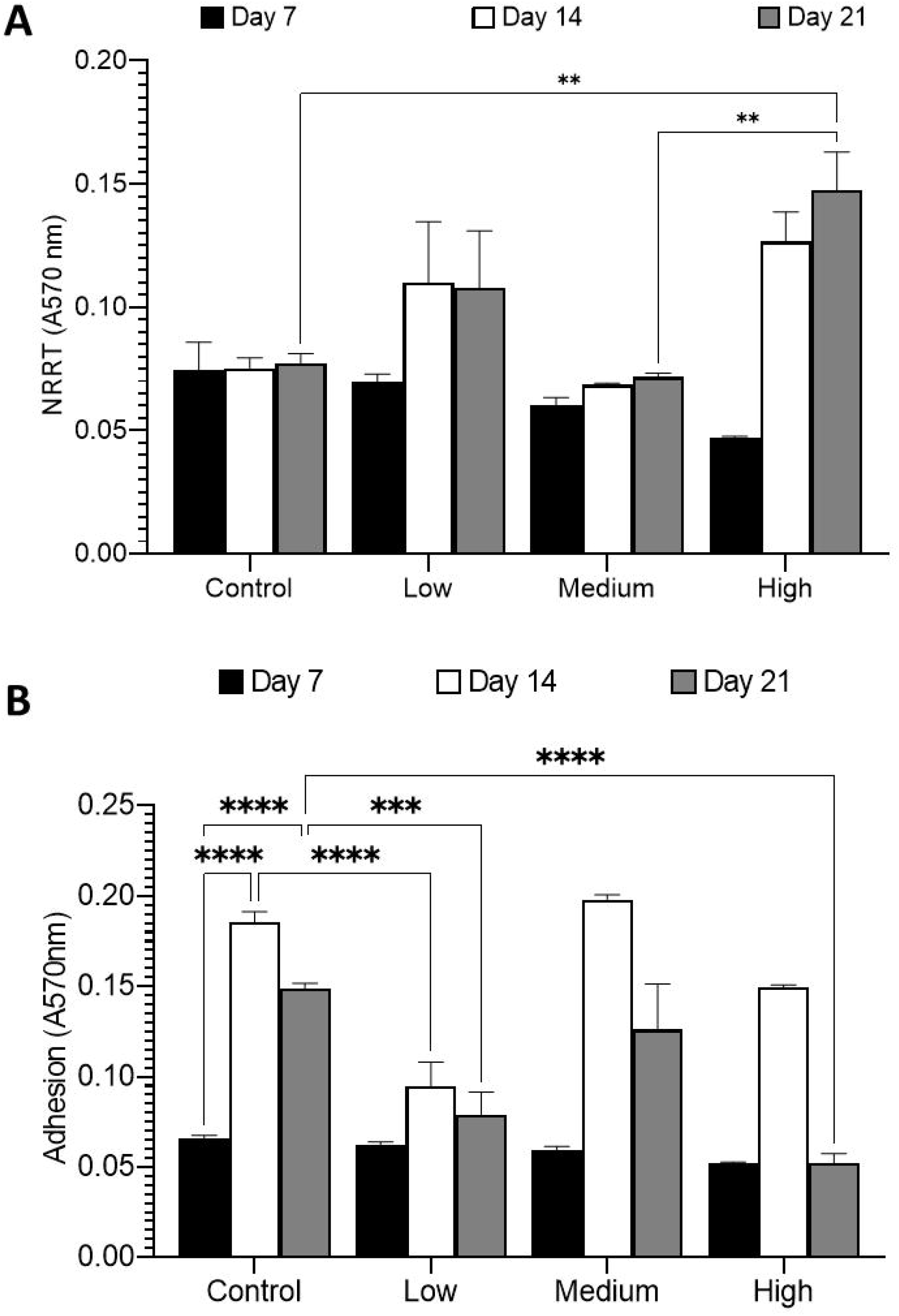
Membrane related events affected by ionising radiation in *Galleria mellonella. G. mellonella* larvae were exposed to gamma radiation from a Cs-137 source for up to 21 days and at the indicated times, pooled haemolymph was measured for neutral red retention **(A)**, or adhesion on plastic, spectrophotometrically **(B)**. Significance was determined by two-way ANOVA, and a Tukey’s multiple comparisons test. (****) p ≤ 0.0001, (***) p ≤ 0.001, (**) p ≤ 0.01. Results are expressed as the mean ± SD of at least two independent experiments (n = 36).

Cell adhesion occurs during cell-cell interactions or cell-extracellular matrix interactions via cell-adhesion molecules. This is critical cell function facilitates signal transduction between cells in response to changes in its surroundings. This can take place during cell migration, tissue development, host-pathogen interaction. In humans, changes to cell adhesion can disrupt cellular functions and lead to a variety of diseases, including cancer, arthritis and infections (Harjunpää et al, 2019; Janiszewska et al, 2020; Johansson et al, 1999). Interestingly, with non-irradiated control conditions, there was a significant increase in cell adhesion to plastic from larvae irradiated to 14 (0.185±0.006, *p* < 0.0001) or 21 (0.149±0.003, *p* < 0.0001) days compared to 7 days (0.066±0.002). At low radiation, adhesion was significantly reduced at the 14- and 21-day time point compared to the non-radiated control. Adhesion was also significantly reduced at high radiation dose at 21 days (0.052±0.005 c.f. control = 0.149±0.003, *p* < 0.0001; **Figure 9B**). This suggests radiation stress negatively – albeit inconsistently – affects haemocyte adhesion to plastic surfaces.

### Radiation affects cellular function *in vitro*

Finally, we wanted to ascertain if some of these cellular functions were <u>directly</u> affected by radiation and not due to radiation-induced bystander effects (Zhou et al, 2005). Therefore, haemocytes were taken from larvae *ex vivo*, cultured for 48 hr *in vitro* and melanisation and pinocytosis determined. Firstly, we found only cultured haemocytes exposed to low dose radiation produced significantly more melanin than its control (0.107±0.008 c.f. control = 0.078±0.004, *p* = 0.003; **Figure 10A**). Interestingly, this result is similar to the whole larvae irradiated for 28 days (**Figure 8B**), suggesting a form of radioprotection triggered by the *G. mellonella* larvae for up to 28 days.

**Figure 10.**
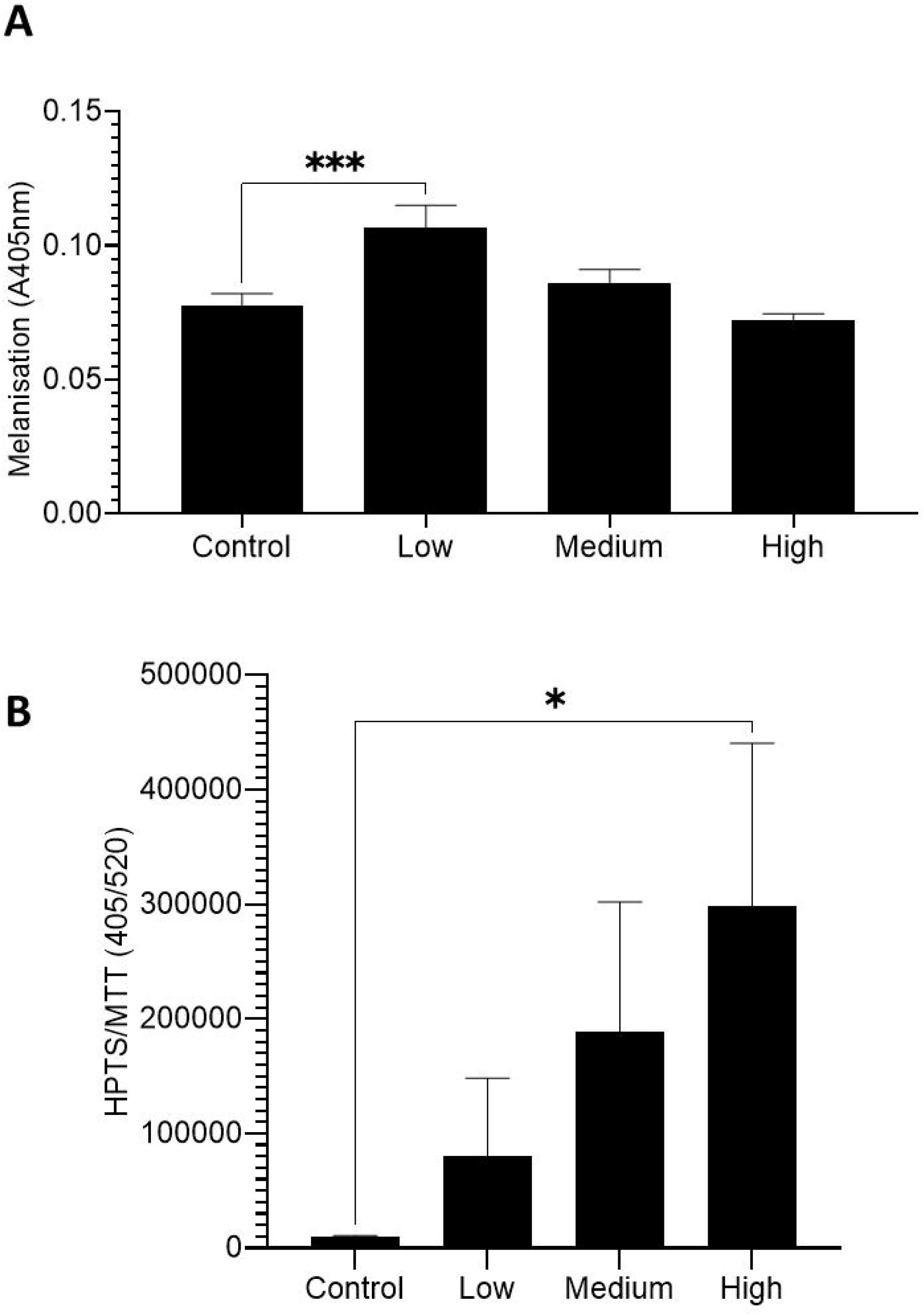
Radiation affected melanin production and endocytosis with cells from *Galleria mellonella*. Haemocytes pooled from *G. mellonella* larvae were cultured and exposed to gamma radiation from a Cs-137 source for 48 hours and measured for melanisation **(A)**, or cell viability and endocytosis, spectrophotometrically **(B)**. Significance was determined by two-way ANOVA, and a Tukey’s multiple comparisons test. (***) p ≤ 0.001, (*) p ≤ 0.05. Results are expressed as the mean ± SD of at least three independent experiments (n = 16).

Furthermore, we explored pinocytosis which is a non-selective fluid-phase endocytosis via the plasma membrane. This process is critical for normal cell function and homeostasis, by sampling the surrounding environment in order to regulate signal processes for adhesion/motility or antigen uptake, etc. A simple method of studying this is through the use of fluid-phase markers such as HPTS (Bugarcic et al, 2012). We found there was a dose dependent uptake of HPTS, with no significant difference with haemocyte viability (data not shown). Haemocytes exposed to high radiation dose showed significantly higher fluid uptake compared to other samples and the non-irradiated control (297884.6±142625.4 c.f. control = 9412.4±1417.5, *p* = 0.02; **Figure 10B**). This suggests radiation actively promotes pinocytosis in haemocytes.

## DISCUSSION

Ionising radiation is known to cause genetic changes, which can have both stimulatory and inhibitory effects on growth and development in all living organisms. In this study, we examined the dose dependent impact of chronic low dose gamma radiation has on the immunobiology of the ecologically and biomedically important insect, *Galleria mellonella*. Understanding the effects of low dose radiation can have significant impact on nuclear material and environmental management, agriculture and pest management. The University of Stirling has a climate-controlled radiation facility that mimics the radiation conditions of the Chernobyl Exclusion Zone through the use of a gamma-emitting caesium-137 source (Beresford et al, 2020). By placing the larvae at different distances from the caesium-137 source and verifying the dose rates during the experiments using dosimeters, we can confirm the dose received at each position. We noticed that, while there was no discernible weight change over a long period of time from irradiated larvae, it is likely due to the “age” of *G. mellonella* larvae which, in this study, were final instars that tend not to feed (Nielsen and Brister 1979; Kwadha et al 2017). This lack of feeding differs from another lepidopteran larvae, the silkworm (*Bombyx mori*), which still feeds across all instars, especially at the late instars (Tayade and Jawale 1984; Radhakrishnan and Periasamy 1986; Bahar et al 2011).

Interestingly, at higher temperature (25 °C), there was a similar response with pupation as it was faster with larvae exposed to a higher dose of radiation and likely due to a response to environmental stressors as seen with silkworms exposed to non-ionising (ultraviolet) radiation (Nojima et al 2019). Incidentally, *G. mellonella* fed an altered diet also triggered early pupation (Emery et al 2021; reviewed by Zhang et al., 2019) and this was observed in other insects exposed to environmental stressors such as house fly (exposed to ozone; Levy et al, 1972) and butterfly species (exposed to haze smoke; Tan et al, 2018) – though those stressors have other significant impacts on the survivability and development of those insect models, including smaller pupae and emergent adults. Interestingly, it is the faecal load produced by *G. mellonella* exposed to ionising radiation that produced a significant effect. Therefore, while there were higher levels of viable bacteria and faecal pellets from larvae exposed to medium and high dose of radiation at the early earlier time points, these were lowered over later time points. This suggests an initial level of environmental stress placed on the larvae, which over time, the insect acclimates to. This effect of environmental stress leading to increased faecal pellet production can also be observed in Mongolian gerbils (Barone et al, 1990; Okano et al, 2005). It will not be surprising if the nature of viable bacteria varies under ionising radiation as it is known that stress (be it psychological, physical and environmental) alters the gut microbiota (Karl et al, 2018; Emery et al, 2021). For example, ozone (O_3_) and nitrogen dioxide (NO_2_) as air pollutants alter the human gut microbiome (Fouladi et al, 2020).

Furthermore, pupation was faster with *G. mellonella* larvae exposed to a higher dose of radiation. We suggest that this accelerated life cycle of the wax moth is due to a physiological response to environmental stress. Insects respond differently to environmental stress and pollution with populations either increasing or decreasing in response to changes to climate or the presence of particular pollutants (reviewed by Pimentel, 1994). Frazier et al. (2006) reported that the population of insects was predicted to increase with the increase in temperature in the environment, which confirms that environmental stressors can alter the physiology of insects and is not necessarily detrimental to the life and wellbeing of insects in our environment. However, alterations to cellular processes need to be further explored as it is not well studied.

Like humans, *G. mellonella* produces melanin in response to radiation. Exposure of *G. mellonella* larvae to ionising radiation alters its immunity, so they overproduce melanin and capsule in a (radiation) dose dependent manner which decreases over time. This priming effect is complemented by a dose-independent increase of haemocyte counts over time. The overall short-term priming effect of radiation on the larvae is it allows replication of pathogens presumably within encapsulation thereby avoid triggering a full immune response with the increase in haemocyte counts. The long-term effect of radiation on larvae is its susceptibility to infection. At higher radiation dose, the immune system might be compromised which could explain the slight increase in fungal load, at least for one species. Interestingly, *ex vivo* haemocytes cultured *in vitro* showed sensitivity to low dose radiation as evidenced from the production of melanin. This is likely due to the lack of the exoskeleton and components of the humoral response involved with melanin production e.g., pro-phenoloxidases (PPOs) (Whitten and Coates, 2017). This corroborates a previous study where *G. mellonella* larvae exposed to 100 – 400 Gy γ-radiation had altered haemocyte morphology, with cytoplasmic vacuolisation and distended mitochondria, as observed in transmission electron micrographs (El-Kholy et al, 2010). In insects, in an open body cavity, overproduction of melanin in response to a microbial infection can become highly toxic, and therefore there are trade-offs between front-loading with melanin and the clearance of pathogens (Lim et al, 2018; Krachler et al, 2021; Smith et al, 2021). Furthermore, any chemical or stressor that reduces melanin allows pathogens to proliferate and cause infection. For example, the herbicide, glyphosate decreases the size of melanised nodules (capsule), thereby allowing proliferation of the yeast *Cryptococcus neoformans*. Interestingly, glyphosate’s mechanism of melanin inhibition involves disruption of reaction of species (ROS) production required for the production of melanin (Smith et al, 2021).

Another area of cell damage due to ionising radiation is through lysosomes, which are important subcellular organelles that contain enzymes that carry out protein degradation and detoxify some foreign compounds (reviewed by Bouhamdani et al, 2021). Lysosomal formation merge with other vesicles involved with digestive pathways such as phagocytosis, endocytosis and autophagy. Therefore the lysosomal membrane must remain intact to protect the cell and any damage to its integrity will lead to release of enzymes or engulfed toxic compounds which leads to oxidative damage and cell death (Moore et al., 2007, 2009a, 2009b). Intra-lysosomal retention of neutral red is a measure of the stability of the lysosomal membrane and therefore a useful indicator of the health of the cell as exposure to stressors leads to compromised lysosomal membrane (Borenfreund and Puerner, 1985; Moore et al., 2009; Piola et al., 2013; Hu et al, 2015). It was reported that 0.1/2.0 Gy of ionising radiation (IR from X-rays) induced significant lysosomal permeability with an increase in ROS in directly irradiated cells, in contrast with a decrease in ROS in bystander cells (Bright et al 2015). That conclusion complements with the lysosomal and total haemocyte count experiments conducted in this study – which showed that the destabilisation of lysosomal membrane at chronic exposure of high dose radiation had a cumulative effect which corresponded with decreased total haemocyte counts. Furthermore, lysosome exocytosis delivers adhesion-related molecules and therefore, lysosomal damage with the release of ROS impacts cell adhesion (Chiarugi et al, 2003; reviewed by Pu et al, 2013). We found – in non-radiated control – baseline adhesion increases, possibly due to release of ROS from vesicles due to the relative (older) age of cells, as reported elsewhere (Hagen et al, 1997; Lee et al, 1999). Therefore, ionising radiation destabilises lysosomal membranes, reduces adhesion and other adhesion-related events such as endocytosis (Cougoule et al, 2004) as evidenced with the *in vitro* HPTS assays.

Therefore, our study demonstrates an interplay between the immune responses mounted by the insect during irradiation and the damage (or not) produced by the pathogen. Alteration to the activation of immune responses can be detrimental to the host, and as such, pathogenicity is a function of both microbial traits and host’s immune response. This integrated view of pathogenicity in insects termed the “damage threshold hypothesis” which is reminiscent of the “damage response framework” observed in mammals (Moreno-García et al, 2014; Casadevall and Pirofski, 2003). This study suggests *G. mellonella* larvae is not only suitable as a tractable model organism to study cellular function exposed to ionising radiation, but also to better probe the full spectrum of microbe–host interactions in insects.

## ACKNOWLEDGEMENTS

JL would like to thank his family for their help and support during the writing of this manuscript and acknowledge the University of Stirling and the Carnegie Trust for the Universities of Scotland (RIG008296) for some funds. DC acknowledges the funding for the radiation facility as part of the TREE (Transfer-Exposure-Effects) consortium under the RATE programme (Radioactivity and the Environment), funded by NERC, the Environment Agency and Radioactive Waste Management Ltd (NE/L000369/1). CJC would like to acknowledge Susan and Lonán Coates for their support.

