## Supplementary figures and images for "Immune-modulatory effects of low dose γ-radiation on wax moth (*Galleria mellonella*) larvae"

### Supplementary Figure 1

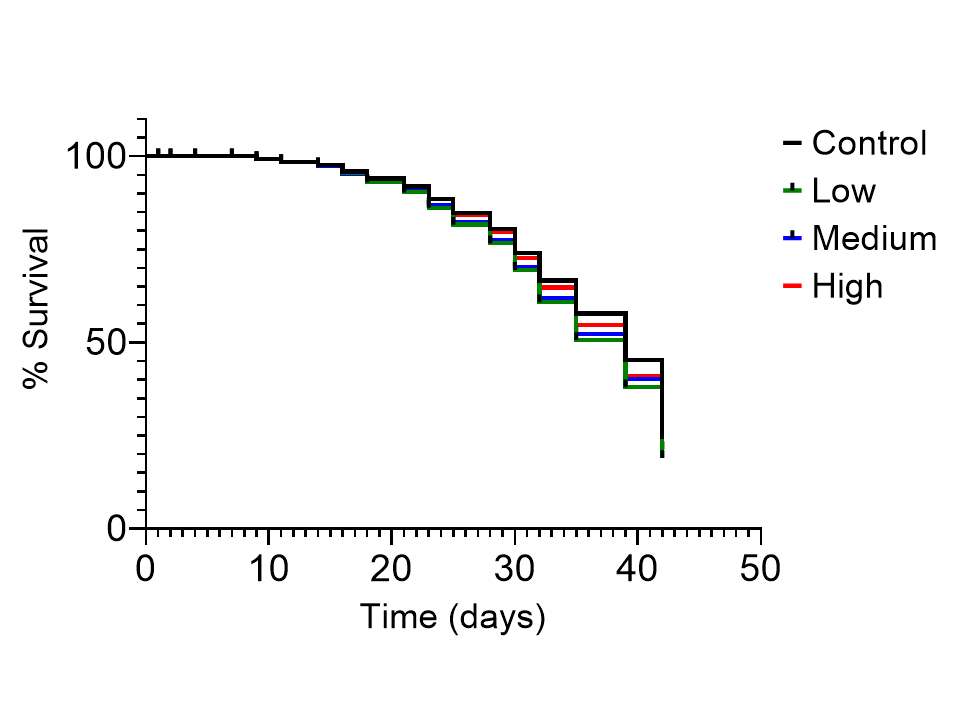
